# Destructive disinfection of infected brood prevents systemic disease spread in ant colonies

**DOI:** 10.1101/116657

**Authors:** Christopher D. Pull, Line V. Ugelvig, Florian Wiesenhofer, Simon Tragust, Thomas Schmitt, Mark J.F. Brown, Sylvia Cremer

## Abstract

Social insects protect their colonies from infectious disease through collective defences that result in social immunity. In ants, workers first try to prevent infection of colony members. Here, we show that if this fails and a pathogen establishes an infection, ants employ an efficient multicomponent behaviour – “destructive disinfection” – to prevent further spread of disease through the colony. Ants specifically target infected pupae during the pathogen’s non-contagious incubation period, relying on chemical ‘sickness cues’ emitted by pupae. They then remove the pupal cocoon, perforate its cuticle and administer antimicrobial poison, which enters the body and prevents pathogen replication from the inside out. Like the immune system of a body that specifically targets and eliminates infected cells, this social immunity measure sacrifices infected brood to stop the pathogen completing its lifecycle, thus protecting the rest of the colony. Hence, the same principles of disease defence apply at different levels of biological organisation.

## Introduction

Pathogen replication and transmission from infectious to susceptible hosts is key to the success of contagious diseases [1]. Social animals are therefore expected to experience a greater risk of disease outbreaks than solitary species, because their higher number of within-group interactions will promote pathogen spread [2–4]. As a consequence, traits that mitigate this cost should have been selected for in group-living animals as an essential adaptation to social life [5,6].

Eusocial insects (termites, ants and the social bees and wasps) live in complex societies that are ecologically successful and diverse. They are typically single-family colonies comprising one or a few reproducing queens and many sterile workers. Both of these castes are highly interdependent: the queens are morphologically specialised for reproduction and cannot survive without the assistance of the workers; conversely, the workers cannot reproduce, but gain inclusive fitness by raising the queen’s offspring [7]. Consequently, social insects societies have become single reproductive units, where natural selection acts on the colony instead of its individual members [8,9]. This has parallels to the evolution of complex multicellular organisms, where sterile somatic tissue and germ line cells form a single reproducing body. Hence, social insect colonies are often termed “superorganisms” and their emergence is considered a major evolutionary transition [8–11]. Since evolution favours the survival of the colony over its members, selection has resulted in a plethora of cooperative and altruistic traits that workers perform to protect the colony from harm [5,8,12,13]. In particular, social insects have evolved physiological and behavioural adaptations that limit the colony-level impact of infectious diseases, which could otherwise spread easily due to the intimate social interactions between colony members [12,14–16]. These defences are performed collectively by the workers and form a layer of protection known as “social immunity” that, like the immune system of a body, protects the colony from invading pathogens [12,17].

Our understanding of how social immunity functions is based mostly on the first line of defence that reduces the probability of pathogen exposure and infection. It is well known for example that social insects avoid pathogens, like fungal spores, in their environment, and perform sanitary care when nestmates come into contact with them [18–22]. In ants, sanitary care involves grooming and the use of antimicrobial secretions to mechanically remove and chemically disinfect the pathogen, reducing the likelihood that pathogen exposure leads to the development of an infection [21,22]. However, what happens when sanitary care fails and a pathogen successfully infects an ant, with the consequent potential to create an epidemic, remains poorly understood. In a body, infected cells are eliminated by the immune system to prevent the proliferation and systemic spread of pathogens through the tissue. Since infected ants become highly contagious to their nestmates [23,24], we hypothesised that they should have evolved an analogous mechanism to detect and contain lethal infections in individuals as early as possible, to prevent disease outbreaks in the colony.

To test this hypothesis, we exposed pupae of the invasive garden ant, *Lasius neglectus*, to a generalist fungal pathogen, *Metarhizium brunneum*. When the infectious conidiospores of this fungus come into contact with insect cuticle, they attach, germinate and penetrate the host cuticle within 48 h to cause internal infections. After a short, non-infectious incubation period of a few days, a successful fungal infection then induces host death, after which the fungus replicates and releases millions of new infectious conidiospores in a process called sporulation [23,25]. Previous work found that brood infected with *Metarhizium* is removed from the brood chamber, however, it is unknown how the ants then respond to the infection [26,27]. Here we demonstrate that ants detect infected pupae during the pathogen’s non-infectious incubation period and react by performing a multicomponent behaviour. To investigate this response we used a series of behavioural and chemical experiments to determine its function and underlying mechanisms. Finally, we tested the impact of the multicomponent behaviour on the pathogen’s ability to complete its lifecycle and cause a systemic colony infection.

## Results

### Destructive disinfection of lethally infected pupae

We exposed ant pupae to one of either three dosages of *Metarhizium* conidiospores or a sham control. We observed that ants tending pathogen-exposed pupae prematurely removed the pupae from their cocoons in a behaviour we termed “unpacking”, whereas control pupae were left cocooned (Fig 1A-B, Video S1; Cox proportional hazards regression: likelihood ratio test (LR) χ^2^ = 55.48, df = 3, *P* = 0.001; post hoc comparisons: control vs. low, *P* = 0.004; low vs. medium, *P* = 0.006; medium vs. high = 0.024; all others, *P* = 0.001). Unpacking occurred between 2-10 d after pathogen exposure, but sooner and more frequently at higher conidiospore dosages (Fig 1B). As unpacking was a belated response to pathogen exposure and we were unable to remove any conidiospores from the cocoon or the unpacked pupae (Fig S1), we concluded that the ants were not performing unpacking to simply dispose of the contaminated cocoons. Instead, we postulated that unpacking was a response to successful infection. At the time of unpacking, the majority of pupae were still alive (Fig S2) and fungal outgrowth had not yet occurred (Fig 1F). Hence, to test if the ants were reacting to early-stage infections, we removed both unpacked and non-unpacked pathogen-exposed cocooned pupae from the ants and incubated them under optimal conditions for fungal outgrowth. We found that, on average across the conidiospore dosages, 90% of unpacked pupae harboured infections that sporulated in the absence of the ants. In contrast, only 25% of non-unpacked pupae were infected (generalised linear model [GLM]: overall LR χ^2^ = 21.52, df = 3, *P* = 0.001; cocooned vs. unpacked pupae: LR χ^2^ = 18.5, df = 1, *P* = 0.001; conidiospore dose: LR χ^2^ = 0.42, df = 2, *P* = 0.81). We therefore concluded that the ants were detecting and unpacking pupae with lethal infections during the asymptomatic incubation period of the pathogen’s lifecycle. At this time point the fungus is non-infectious and so there is no risk of the ants contracting the disease.

Next, we filmed ants presented with pathogen-exposed pupae and compared their behaviour before and after unpacking. Prior to unpacking, we observed the typical sanitary care behaviours reported in previous studies [20,22,23,28]. Namely, the ants groomed the pupae (Fig 1C), which has the dual function of removing the conidiospores and applying the ants’ antimicrobial poison [22]. In *L. neglectus*, the poison is mostly formic acid and is emitted from the acidopore at the abdominal tip, where the ants actively suck it up and transiently store it in their mouths until application during grooming. Additionally, the ants can spray their poison directly from the acidopore; yet, this behaviour is rarely expressed during sanitary care (about once every 28 h; Fig 1D) [22]. However, after unpacking, we observed a set of behaviours markedly different to sanitary care (Fig 1A, Video S1). The ants sprayed the pupae with poison from their acidopore approx. 15-times more frequently than during sanitary care (∼ 13-times/d; Fig 1D; generalised linear mixed model [GLMM]: LR χ^2^ = 17.04, df = 1, *P* = 0.001), and increased grooming by 50% (Fig 1C; linear mixed effects regression [LMER]: LR χ^2^ = 145.26, df = 1, *P* = 0.001). Given that there was no fungus to remove at the time of unpacking, the increase in grooming probably functioned solely to apply poison from the oral store [22]. Furthermore, the ants repeatedly bit the pupae to make perforations in their cuticles (Fig 1E; GLMM: LR χ^2^ = 39.44, df = 1, *P* = 0.001). Together these three behaviours resulted in the death of the pupae and left their corpses heavily damaged and coated in the ants’ poison (Fig 1G, Fig S2, Fig S3). Accordingly, we named the combination of unpacking, grooming, poison spraying and biting “destructive disinfection”, and performed a series of experiments to determine its function.

**Fig 1.**
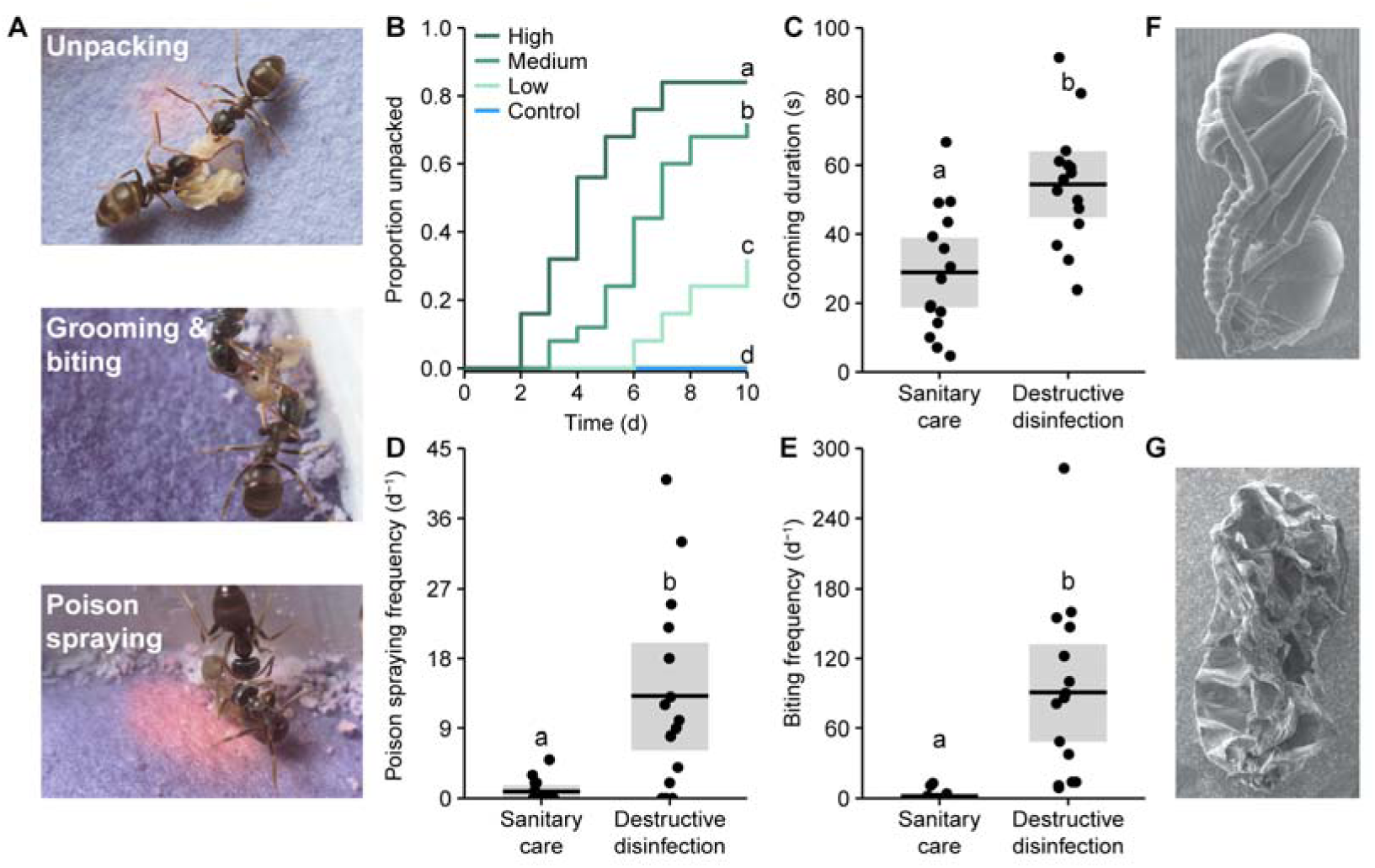
Ants perform destructive disinfection in response to lethal fungal infections of pupae. (A) Destructive disinfection starts with the unpacking of pupae from their cocoons and is followed by grooming, poison spraying and biting (ants housed on blue pH-sensitive paper to visualise acidic poison spraying, which shows up pink). (B) Unpacking occurred significantly more in pupae exposed to fungal conidiospores and was dose-dependent, occurring sooner and in higher amounts as the dose of conidiospores increased (letters denote groups that differ significantly in Tukey post hoc comparisons [*P* < 0.05]). (C-E) Comparison of the ants’ behaviour between sanitary care and destructive disinfection. Destructive disinfection is characterised by increases in grooming duration, poison spraying frequency and biting frequency (all data points displayed; lines ± shaded boxes show mean ± 95% confidence intervals [CI]; letters denote groups that differ significantly in logistic regressions [*P* < 0.05]). (F) Scanning electron micrographs (SEM) of an asymptomatic pupa immediately after unpacking, and (G) of a destructively disinfected pupa 24 h later.

### Chemical detection of internal infections

Firstly, we wanted to know how the ants identify internal infections during the pathogen’s non-contagious incubation period, when pupae were still alive and showed no external signs of disease. As ants use chemical compounds on their cuticles to communicate complex physiological information to nestmates [29], we speculated that infected pupae may produce olfactory sickness cues. We washed infected pupae in pentane solvent to reduce the abundance of their cuticular hydrocarbons (CHCs). When pentane-washed pupae were presented to ants, there was a 72% reduction in unpacking compared to both non- and water-washed infected pupae (Fig 2A; GLM: LR χ^2^ = 12.2, df = 2, *P* = 0.002; Tukey post hoc comparisons: water-washed vs. non-washed, *P* = 0.79; all others, *P* = 0.009). As pentane-washed pupae had lower abundances of CHCs (Fig S4), this result indicates that the ants use one or more cuticle compounds to detect the infections.

Gas chromatography-mass spectrometry (GC–MS) analysis of the solvent wash confirmed that unpacked pupae have distinct chemical profiles compared to non-infected control pupae, whilst cocooned (non-unpacked) pathogen-exposed pupae were intermediate (Fig 3B, Fig S5; perMANOVA: *F* = 1.49, df = 46, *P* = 0.002; post hoc perMANOVA comparisons: unpacked vs. control, *P* = 0.003; unpacked vs. cocooned, *P* = 0. 79; cocooned vs. control, *P* = 0.08). Most chemical messages in social insects are encoded by quantitative shifts of several compounds [29]. Correspondingly, we found that 8 out of the 24 CHCs identified (Table S1) had higher relative abundances on unpacked pupae compared to control pupae (Fig 3C-F, Fig S5; all Kruskal-Wallis [KW] test statistics and post hoc comparisons in Table S2). Moreover, four of these CHCs were also present in relatively higher quantities on unpacked pupae compared to the non-unpacked cocooned pupae. Hence, several specific CHCs probably accumulate on infected pupae over time, eventually reaching an amount that, relative to the other compounds, is sufficient to elicit destructive disinfection. This corresponds to current models of social insect behaviour, where the likelihood of a response depends on stimuli exceeding a certain threshold [30,31]. Interestingly, the four CHCs specifically increased on unpacked pupae were all long-chained CHCs (carbon chain length C_33-35_) with a low volatility, meaning that the ants have to be close to or touching the pupae to detect them [32]. As ants keep pupae in large piles, using low-volatility CHCs may be important so that the ants accurately identify the sick pupae and do not mistakenly destroy healthy ones.

**Fig 2.**
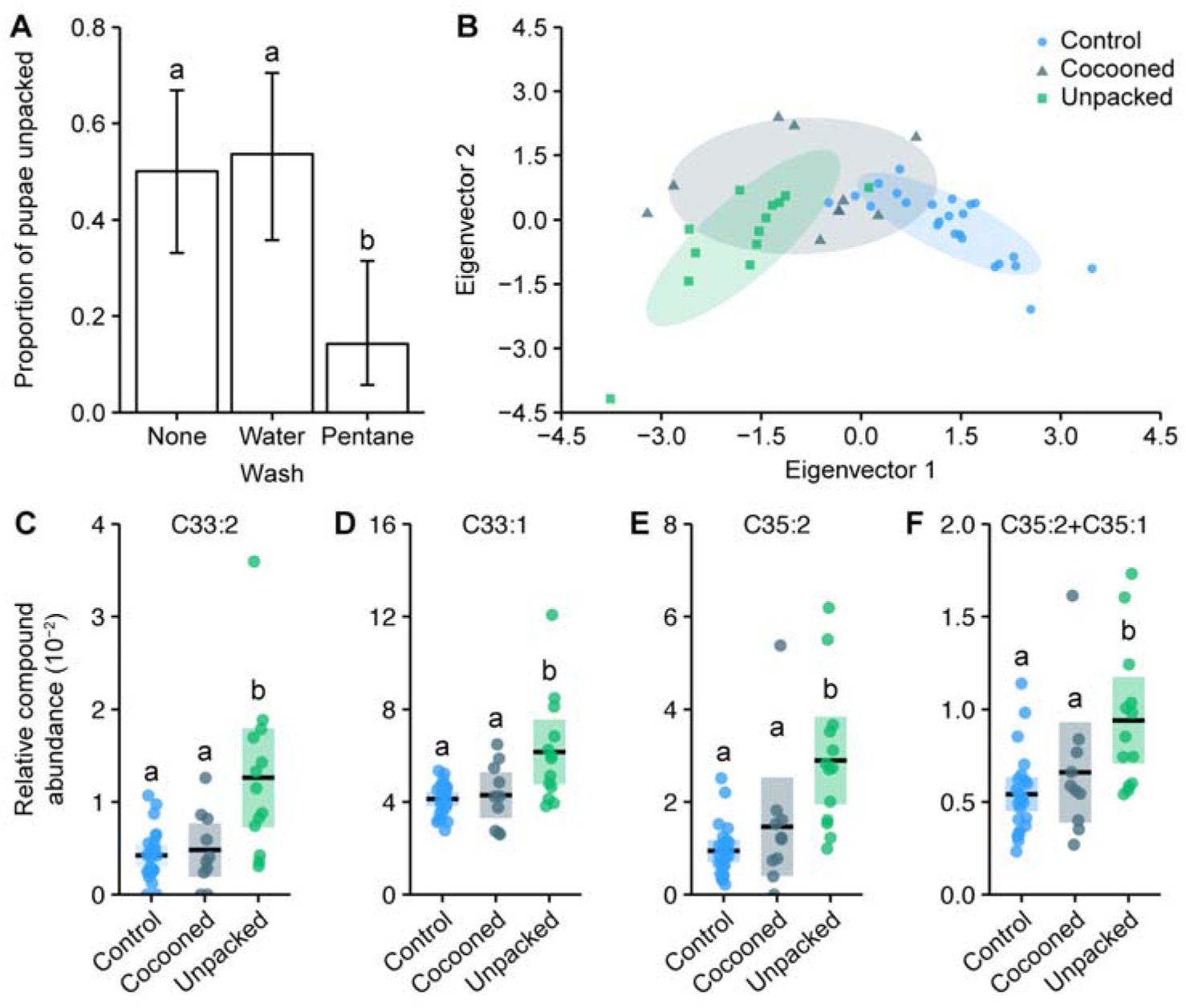
Destructive disinfection is induced by changes in the chemical profile of infected pupae. (A) Pupae washed in pentane solvent to reduce the abundance of their cuticular hydrocarbons (CHCs) were unpacked less than unwashed or water-washed pupae (positive and handling controls, respectively; error bars show ± 95% CI; letters specify significant Tukey post hoc comparisons [*P* < 0.05]). (B) Unpacked pathogen-exposed pupae have distinct chemical profiles compared to sham-treated control pupae. Pathogen-exposed pupae that were not unpacked (cocooned group) have intermediate profiles (axes show discriminant analysis of principle components eigenvectors). (C-F) The four CHCs with higher relative abundances on unpacked pupae compared to both control and cocooned pupae, (C) Tritriacontadiene, C33:2 (D), Tritriacontene, C33:1 (E), Pentatriacontadiene, C35:2 (F) co-eluting Pentatriacontadiene and Pentatriacontene, C35:2+C35:1 (all data points displayed; line ± shaded box show mean ± 95% CI; letters specify groups that differ significantly in KW test post hoc comparisons [*P* < 0.05]).

### Destructive disinfection prevents pathogen replication

We next tested if destructive disinfection prevents pupal infections from replicating and becoming infectious. Pathogen-exposed pupae were kept with groups of ants (8 ants per pupae per group) until unpacking. They were then left with the ants for a further 1 or 5 d before being removed and incubated for fungal growth. We compared the number that subsequently sporulated to pathogen-exposed pupae kept without ants. Whilst 90% of pupae contract infections, destructive disinfection significantly reduced the proportion of pupae that sporulated and hence became infectious (Fig 3A; GLM: LR χ^2^ = 40.47, df = 2, *P* = 0.001; Tukey post hoc comparisons: 1 vs. 5 d, *P* = 0.04; all others, *P* = 0.001). After only 1 d, the number of destructively disinfected pupae that sporulated decreased by 65%. With more time, the ants could reduce the number of pupae sporulating even further by 95%. Since the pupae were removed from the ants for fungal incubation, we can conclude that destructive disinfection permanently prevents pathogen replication. We repeated this experiment with a smaller number of ants (3 ants per pupae per group) to investigate how group size influences the success of destructive disinfection. Smaller groups of ants were less efficient than larger ones: although they could still inhibit > 90% of pupal infections within 5 d of unpacking, pupae tested for infection after 1 d still sporulated 70% of the time (Fig S6; GLM: LR χ^2^ = 35.23, *P* = 0.001; Tukey post hoc comparisons: 0 vs. 1 d, *P* = 0.2; 0 vs. 5 d, *P* = 0.001; 1 vs. 5 d, *P* = 0.002). As the effectiveness of destructive disinfection increased with the amount of time the ants had, as well as with the number of ants present, we inferred that there must be a limiting factor affecting the inhibition the pathogen.

To study the underlying mechanisms of destructive disinfection, we performed its different components – unpacking, biting and poison spraying – *in vitro* to test for their relative importance and potential synergistic effects. We simulated unpacking by removing the cocoons of the pupae manually, and the cuticle damage caused by biting using forceps. Previous work establishing the composition of *L. neglectus* poison [22] allowed us to create a synthetic version for use in this experiment (60% formic acid and 2% acetic acid, in water; applied at a dose equivalent to what ants apply during destructive disinfection; Fig S8), with water as a sham control. We then performed these ‘behaviours’ in different combinations in a full-factorial experiment. We found that all three behaviours must be performed in the correct order and interact to prevent pathogen replication (overview graph showing odds ratios of sporulation in Fig 3B, full data dataset displayed in Fig S7; GLM: overall LR χ^2^ = 79.9, df = 5, *P* = 0.001; interaction between behaviours LR χ^2^ = 20.6, df = 2, *P* = 0.001; all post hoc comparisons in Table S3). As in sanitary care, the poison was the active antimicrobial compound that inhibited fungal growth (Fig S7, Table S3 [21,22]). However, for the poison to function the pupae had to be removed from their cocoons and their cuticles damaged. Firstly, this is because the cocoon itself is hydrophobic and thus prevents the aqueous poison from reaching the pupae inside (Fig S9). Secondly, as the infection is growing internally at the time of unpacking, the cuticle must be broken in order for the poison to enter the hemocoel of the pupae. This is achieved with the perforations created by the ants biting the pupal cuticle. As the active antimicrobial component, we concluded that the poison is probably the limiting factor determining whether destructive disinfection is successful. Because the poison has a slow biosynthesis and each ant can only store a limited amount [22,33], it would explain why destructive disinfection was more likely to be successful the longer the ants had to treat the pupae, and as the number of ants increased (Fig 3A, Fig S6). By sharing the task of poison synthesis and application, the ants probably increase their chances of preventing the pathogen becoming infectious.

**Fig 3.**
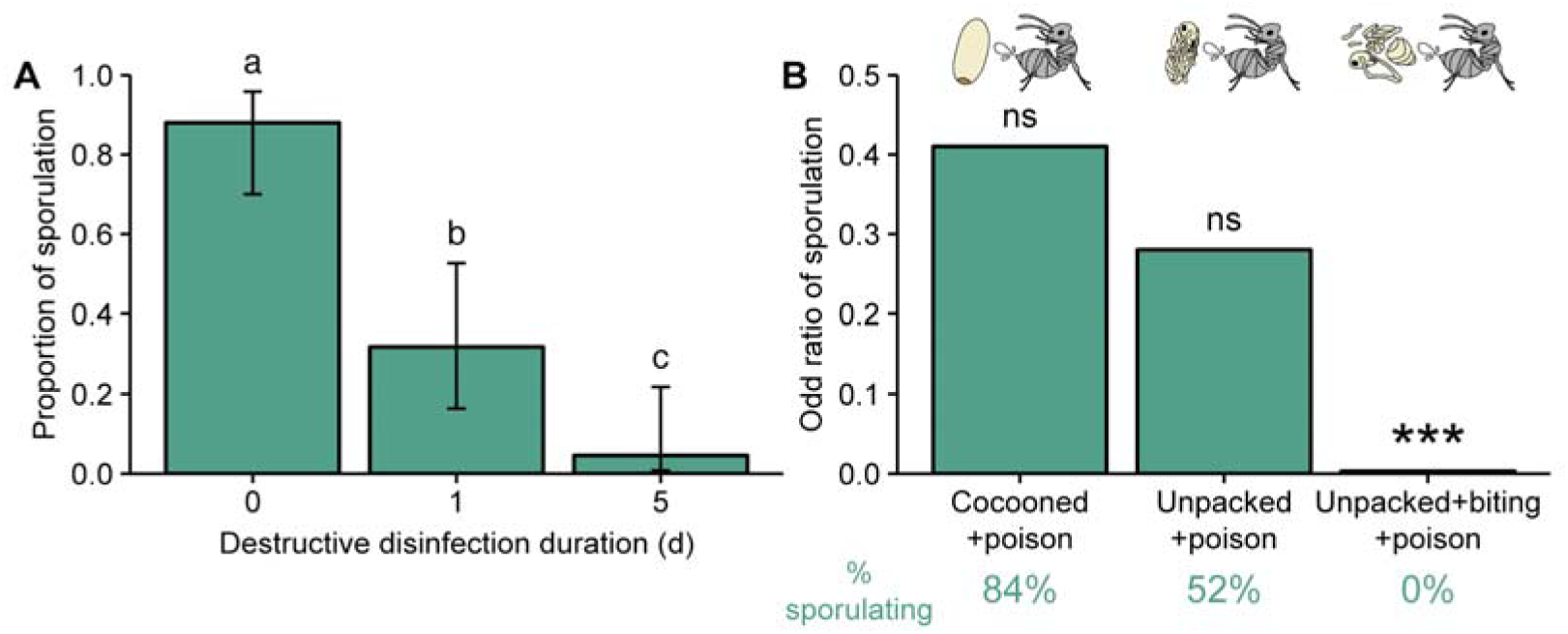
Destructive disinfection by ants prevents pathogen replication. (A) Destructive disinfection greatly reduced the probability of pupae sporulating compared to pupae that received no destructive disinfection (time point 0), and its effectiveness increased with the length of time ants could perform destructive disinfection (1 vs. 5 d; error bars show ± 95% CI; letters denote groups that differ significantly in Tukey post hoc comparisons [*P* < 0.05]). (B) The individual components of destructive disinfection (unpacking, biting and poison spraying) interacted to inhibit pathogen replication (% of pupae sporulating in each treatment shown under graph in green). The odds of sporulation for cocooned and unpacked pupae treated with poison were not significantly different to those of control pupae (cocooned pupae treated with water). But when unpacking, biting and poison spraying were combined the odds of sporulation were significantly reduced (logistic regression; ns = non-significant deviation from control, *** = ***P*** < 0.001; complete data set of full factorial experiment displayed in Fig S7 and all statistics in Table S3).

### Disruption of the pathogen lifecycle stops disease transmission

Finally, we investigated the impact of destructive disinfection on disease transmission within a social host group. We created mini-nests comprising two chambers and a group of ants (5 ants per group). Into one of the chambers we placed an infectious sporulating pupa – simulating a failure of the ants to detect and destroy the infection – or a pupa that had been destructively disinfected, and was thus non-infectious. The ants groomed, moved around and sprayed both types of corpses with poison. In the case of the sporulating pupae, all conidiospores were removed from the corpse by the ants. As in previous studies, sporulating corpses were highly virulent [23,24] and caused lethal infections that became contagious after host death in 42% of ants (Fig 4A). However, there was no disease transmission from destructively disinfected pupae (Fig 4A; GLM: LR χ^2^ = 31.32, df = 1, *P* = 0.001). We therefore concluded that by preventing the pathogen from completing its lifecycle destructive disinfection stops intra-colony disease transmission (Fig 4B).

**Fig 4.**
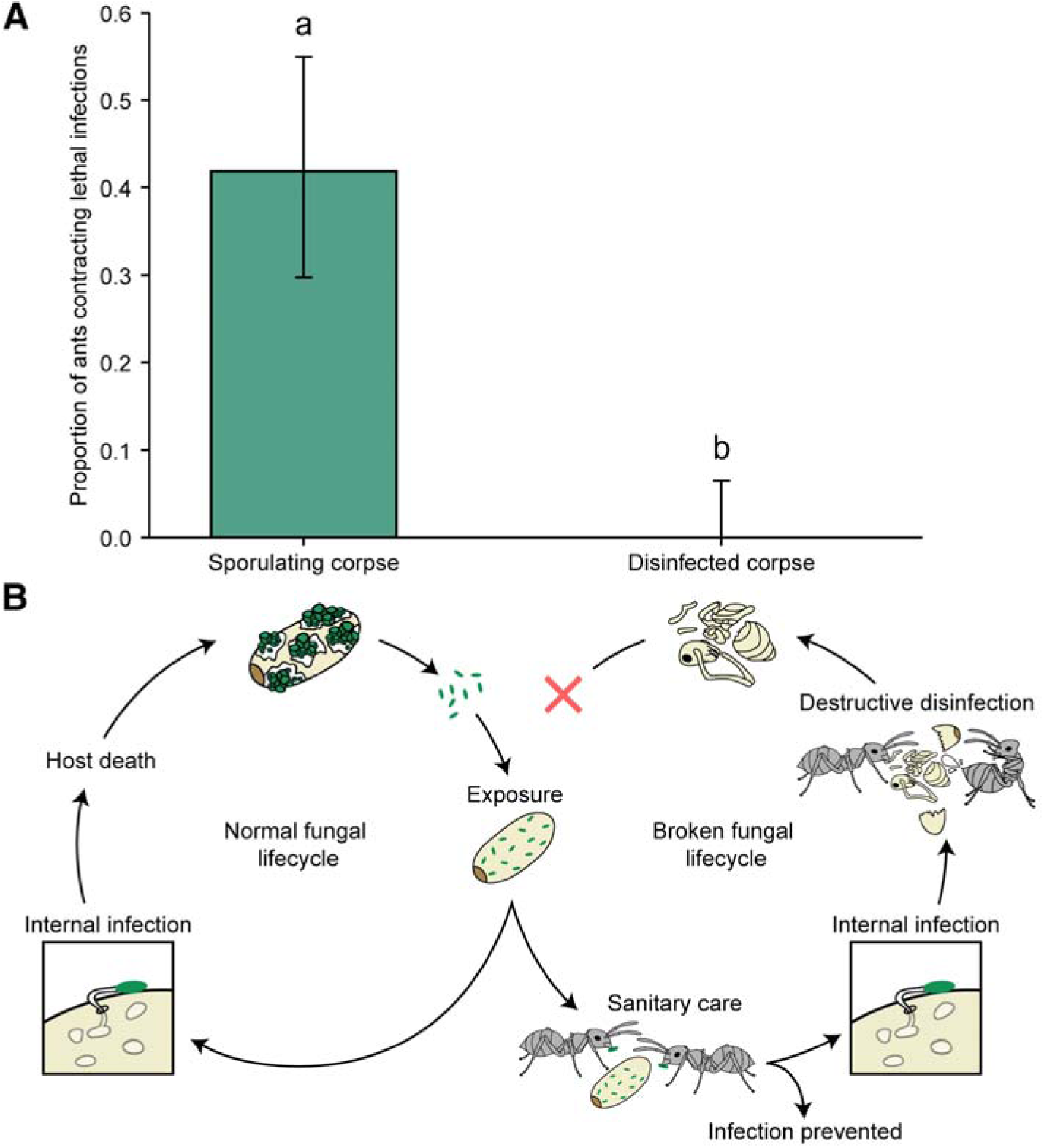
Destructive disinfection stops disease transmission. (A) Ants that interacted with sporulating pupae contracted lethal infections and died from fungal infection in 42% of the cases, whilst there was no disease transmission from destructively disinfected pupae (error bars show ± 95% CI; letters denote groups that differ significantly in a logistic regression [*P* < 0.05]). (B) Overview of normal fungal lifecycle resulting in infectious, sporulating corpses (left) and a broken lifecycle due to the interference of the ants (right). When sanitary care fails to prevent infection in pathogen-exposed individuals, the ants switch to colony-level disease control, i.e. destructive disinfection to stop pathogen replication, resulting in non-infectious corpses.

## Discussion

Ants are extremely hygienic and frequently perform sanitary behaviours to prevent microbial infection of themselves and colony members [12]. However, if these behaviours fail, the colony faces a problem because infections can become highly contagious and cause disease outbreaks [23,24]. In this study, we have characterised a multicomponent behaviour that ants use to fight lethal infections of a common fungal pathogen. Our results show that ants detect infected pupae using chemical signatures whilst the pathogen is still in its non-transmissible incubation period (Fig 2). In contrast to the simple removal of infected brood in honeybees [15], the ants then performed destructive disinfection, utilising their antimicrobial poison for internal disinfection of the host body to stop the pathogen from replicating and completing its lifecycle (Fig 1, Fig 3). Ultimately, this prevented the fungus from infecting new hosts and effectively reduced its fitness to zero (Fig 4). These findings extend our current understanding of collective disease defence in ants, showing that they not only avoid [18], groom [20–22] and isolate pathogens [22,26], but can even interfere with the infectious cycle of the pathogen to actively arrest its establishment and replication in the colonies (Fig 4b). This will have important implications for the evolution of host-pathogen interactions in social insects, as the pathogen is unable to reproduce. More generally, our results reveal the remarkable adaptations that can evolve in superorganisms to avoid disease outbreaks.

We found that destructive disinfection acts like a second line of defence for the colony, when the first, sanitary care, fails to prevent infection. This has parallels to the immune system of the body where defences are layered to prevent pathogen establishment and replication at multiple levels [17]. The first line of defence in the body is made up of mechanical and chemical defences, such as ciliated cells in the lung that move pathogens trapped in mucus out of the body [17]. In ants, grooming and chemical disinfection during sanitary care play an analogous role [20–22]. However, if a pathogen circumvents these defences and a cell is infected, the second line of defence is often a targeted elimination of the cell. This starts with immune cells detecting an infection and then transporting cell death-inducing and antimicrobial compounds into the infected cell by creating pores in its membrane [34–36]. Likewise, our experiments revealed that ants detect sick pupa using chemical compounds on their cuticle. They then unpack the pupa and make perforations in its cuticle, enabling the ants to spray their poison directly into the pupa’s body. In both cases, the second line of defence destroys the infected cell/insect, along with the infection, to prevent transmission [37]. Since the loss of somatic cells and individual insect workers can be tolerated with negligible effects on fitness [17], these analogous strategies are a unique way to clear infections and avoid any further damage to the body and colony, respectively.

Previous studies have suggested that ants might use chemical cues to detect sick colony members, but evidence to support this hypothesis has been lacking [26,38,39]. To our knowledge, we have therefore discovered the first known instance of ants using chemical information to identify and specifically target infected individuals. The chemical compounds with increased abundances on infected pupae are distinct from those that induce the removal of corpses in ants [40–42], and, like in tapeworm-infected ants [43], are not pathogen-derived because they are also present in lower amounts on healthy pupae. This alteration of the hosts’ chemical profile may arise during infection from the breakdown of hydrocarbons by *Metarhizium* penetration [44] or after infection due to an immune response affecting the synthesis of specific hydrocarbons [45,46]. The latter is more likely as the ants only display destructive disinfection once the fungus is growing inside the pupae. Interestingly, two of the four CHCs that were increased on infected pupae also had higher abundances on virus-infected honeybees (Tritriacontadiene [47]) and their brood experiencing a simulated bacterial infection (Tritriacontene [46]). As these compounds belong to the same hydrocarbon substance class – unsaturated hydrocarbons – their common biosynthetic pathway might be upregulated upon infection. This raises the possibility that these hydrocarbons are evolutionarily conserved sickness cues in Hymenopteran social insects. Such cues may have evolved into general sickness signals in social insects as they alert nestmates to the presence of an infection that will harm the colony if it spreads [48]. Similar to the “find-me/eat-me” signals expressed by infected cells in a body [49,50], they will be selected for as they enhance colony fitness by preventing a systemic infection. Hence, altruistic displays of sickness can evolve in superorganisms, even if this results in the destruction of the individual that expresses them.

It is well established that social insects use glandular secretions with antimicrobial properties as external surface disinfectants [51]. However, because these compounds can also harm the host, they should be used with caution inside the colony. For example, the acidic poison *L. neglectus* and other Formicine ants produce is extremely caustic and is used to attack conspecifics [22,33,52]. During sanitary care they apply this poison via grooming because it is probably more accurate and less wasteful than spraying [22]. Moreover, as pathogen-exposed insects typically survive when they receive sanitary care [20–23], conservatively applying the poison may also reduce the damage it causes to individuals that can then continue contributing to the colony. This is supported by our observation that *L. neglectus* will apply large quantities of poison onto pupae only when they become infected. Remarkably, we found that, in addition to being external disinfectants, ants use antimicrobial secretions as internal disinfectants against infections within the bodies of nestmates. Since infected pupae are moribund there is no risk that the ants’ poison is harming individuals with a future role in the colony. Taken together, these observations suggest that ants adjust their behaviours in response to the risk presented to the colony. It would be interesting to explore further how social immunity defences are regulated to prevent collateral damage, or ‘social immunopathology’, within the colony.

Our experiments show that destructive disinfection was highly effective and prevented 95% of infections becoming transmissible. Destructive disinfection will thus keep the average number of secondary infections caused by an initial infection small and the disease will die out within the colony [3]. This may explain why infections of *Metarhizium* and other generalist entomopathogenic fungi like *Beauveria*, though common in the field [53–56], do not seem to cause colony-wide epidemics in ants, but are more numerous in solitary species that lack social immunity [57–59]. Behaviours like destructive disinfection that are able to reduce pathogen fitness to zero could have selected for host manipulation in fungi that specialise on infecting ants, e.g. *Ophiocordyceps* and *Pandora* [60-62]. These fungi force their ant hosts to leave the nest and climb plant stems near foraging trails. There they die and become infectious, releasing new spores that infect ants foraging below. However, ants infected with *Ophiocordyceps* that were experimentally placed back into the nest disappeared [61]. Our study suggests that these ants could have been eliminated through destructive disinfection. Consequently, ant-specialist fungi like *Ophiocordyceps* and *Pandora* may have evolved host manipulation as a means to complete their lifecycle outside of the nest and avoid destructive disinfection [61,62]. In contrast to specialists, generalist pathogens like *Metarhizium* infect a broad range of solitary and social hosts, making it less likely that they evolve strategies to escape social immunity defences [63]. Future work that investigates how social immunity disrupts typical host-pathogen dynamics will shed light on the co-evolution of pathogens and their social hosts [3].

Destructive disinfection has probably evolved in ants because the removal of corpses from the colony alone does not guarantee that disease transmission is prevented [61]. This is because ants place corpses onto midden (trash) sites that are located inside or outside near the nest and regularly visited by midden workers [64–66]. Consequently, midden sites represent a potential source for disease transmission back into the colony. In contrast to ants, honeybees have no middens and corpses are dumped randomly outside of the hive [15]. But because honeybees forage on the wing, it is unlikely that corpses are re-encountered and so removal is sufficient to prevent disease transmission [67]. Termites on the other hand perform a different behaviour, whereby the dead are cannibalised [19,68]. Cannibalism is effective because the termite gut neutralises ingested pathogens [69–71] and has likely evolved because dead nestmates are a source of valuable nitrogen in their cellulose-base diet [72]. The same selective pressure has driven this suite of independently evolved innovations – the need to eliminate or remove infected individuals early in the infectious cycle – with the ants expressing a particularly complex behavioural repertoire. This seems to be a general principle in disease defence as cells are also rapidly detected and destroyed shortly after infection to prevent pathogen spread in multicellular organisms [17]. Understanding how natural selection can result in similar traits at different levels of biological organisation and in organisms with different life histories is a central question in evolutionary biology [8]. Studying the similarities and differences between organismal immunity and social immunity could therefore lead to new insights about how disease defences evolve [17]. For example, the results of our study suggest that equivalent selection pressures can result in convergent defences that protect multicellular organisms and superorganismal insect societies from systemic disease spread. Future work that can link the performance of social immunity defences to colony fitness will therefore provide useful insights into how such traits are selected for over evolutionary time.

## Materials and Methods

### Ant host

We studied the unicolonial invasive garden ant, *Lasius neglectus*, collected in Seva, Spain (41.809000, 2.262194) [55]. Stock colonies were kept at a constant temperature of 23°C with 70% humidity and a day/night cycle of 14/10 h. All experiments were conducted in plastered petri dishes (Ø = 33, 55 or 90 mm) with 10% sucrose solution provided *ad libitum* and environmental conditions were controlled throughout (23°C; 70% RH; 14/10 h light/dark cycles). Care of animals was in accordance with institutional guidelines.

### Fungal pathogen

As a model pathogen, we used the entomopathogenic fungus *Metarhizium brunneum* (strain MA275, KVL 03-143). Multiple aliquots were kept in long-term storage at – 80°C. Prior to each experiment the conidiospores were grown on sabaroud dextrose agar at 23°C until sporulation and harvested by suspending them in 0.05% sterile Triton X-100 (Sigma). The germination rate of conidiospore suspensions was determined before the start of each experiment and was > 90% in all cases.

### Pupal pathogen exposure

Conidiospores were applied in a suspension of 0.05% autoclaved Triton-X 100 at 10^6^ conidia/ml in all experiments unless otherwise stated. Throughout the study, we used cocooned worker pupae of approximately the same age, which was determined by assessing the melanisation of the eyes and cuticle. Single pupae were exposed by gently rolling them in 1 μl of the conidiospore suspension using sterile soft forceps. Pupae were then allowed to air dry for 5-10 min before being used in experiments. This exposure procedure resulted in pupae receiving ∼ 1800 conidiospores, of which 5% (∼ 95 conidiospore) passed through the cocoon and came into contact with the pupa inside (Fig S1).

### Statistical Analysis

Statistical analyses were carried out in R version 3.3.2 [73]. All statistical tests were two-tailed. General(ised) linear and mixed models were compared to null (intercept only) and reduced models (for those with multiple predictors) using Likelihood Ratio (LR) tests to assess the significance of predictors [74]. We controlled for the number of statistical tests performed per experiment to protect against a false discovery rate using the Benjamini-Hochberg procedure (α = 0.05). Moreover, all post hoc analyses were corrected for multiple testing using the Benjamini-Hochberg procedure (α = 0.05) [75,76]. We checked the necessary assumptions of all tests i.e. by viewing histograms of data, plotting the distribution of model residuals, checking for non-proportional hazards, testing for unequal variances, testing for the presence of multicollinearity, testing for over-dispersion, and assessing models for instability and influential observations. For mixed effects modelling, we used the packages ‘lme4’ to fit models [77], ‘influence.ME’ to test assumptions [78], and, for LMERs, ‘lmerTest’ to obtain *P* values [79]. All logistic regressions were performed using either generalised linear models (GLMs) or generalised linear mixed models (GLMMs), which had binomial error terms and logit-link function. The Cox proportional hazards regression was carried out using the ‘coxphf’ package with post hoc comparisons achieved by re-levelling the model and correcting the resulting *P* values [80]. For Kruskal-Wallis (KW) tests and subsequent post hoc comparisons we used the ‘agricolae’ package, which implements the Conover-Iman test for multiple comparisons using rank sums [81]. For the perMANOVA, we used the package ‘vegan’ and performed pairwise perMANOVAs for post hoc comparisons [82]. All other post hoc comparisons were performed using the ‘multcomp’ package [83]. Finally, all graphs were made using the ‘ggplot2’ package [84]. Individual descriptions of statistical analyses are given for all experiments below.

### Unpacking behaviour

To study how ants respond to infections, we exposed pupae to a low (10^4^/ml), medium (10^6^/ml) or high (10^9^/ml) dose of conidiospores or autoclaved Triton X as a sham control (sham control, *n* = 24; all other treatments, *n* = 25). The pupae were then placed into individual petri dishes with two ants and inspected hourly for 10 h/d for 10 d. When the ants unpacked a pupa, it was removed and surface-sterilised [85] to ensure that any fungal outgrowth was the result of internal infections and not residual conidiospores on the cuticle. After sterilisation, we transferred the pupae to a petri dish lined with damp filter paper at 23°C and monitored them for 2 weeks for *Metarhizium* sporulation to confirm the presence of an internal infection (low dose, *n* = 8; medium dose, *n* = 18; high, *n* = 21). In addition, any cocooned pupae that were not unpacked after 10 d were removed from the ants, surface sterilised and observed for sporulation, as above (low dose, *n* = 11; medium dose, *n* = 4; high, *n* = 4). We analysed the effect of treatment on unpacking using a Cox proportional hazards model with Firth’s penalized likelihood, which offers a solution to the monotone likelihood caused by the complete absence of unpacking in the sham control treatment. We followed up this analysis with post hoc comparisons (model factor re-levelling) to test unpacking rates between treatments (Fig 1B). We compared the number of unpacked and cocooned pupae sporulating using a logistic regression, which included pupa type (cocooned, unpacked), conidiospore dose (low, medium, high) and their interaction as main effects. The interaction was non-significant (GLM: LR χ^2^ = 5.0, df = 2, *P* = 0.084); hence, it was removed to gain better estimates of the remaining predictors.

### Images and scanning electron micrographs (SEMs) of destructive disinfection

Photographs of destructive disinfection were captured (Nikon D3200) and aesthetically edited (Adobe Photoshop) to demonstrate the different behaviours (Fig 1A). They were not used in any form of data acquisition. We also made representative SEMs of a pupa directly after unpacking and one after destructive disinfection (24 h after unpacking; Fig 1F-G). As the pupae were frozen at – 80°C until the SEMs were made, we also examined non-frozen pupae taken directly from the stock colony and confirmed that freezing itself does not cause damage to the pupa (not shown).

### Comparison of sanitary care and destructive disinfection behaviours

To observe how the behavioural repertoire of the ants changes between sanitary care and destructive disinfection, we filmed three individually colour-marked ants tending a single pathogen-exposed pupa with a USB microscope camera (Di-Li ® 970-O). To characterise the sanitary care behaviours of the ants, we analysed the first 24 h of the videos following the introduction of the pupa. To study destructive disinfection behaviours, we analysed the 24 h period that immediately followed unpacking. Videos were analysed using the behavioural-logging software JWatcher™ [86]. For each ant (*n* = 15), we recorded the duration of its grooming bouts, the frequency of poison application and the frequency of biting. Grooming duration was analysed using a LMER, having first log-transformed the data to fulfil the assumption of normality (Fig 1C). The frequency of poison application and biting (Fig 1D-E) were analysed using separate GLMMs with Poisson error terms for count data and logit-link function. We included an observation-level random intercept effect to account for over-dispersion in the data [87]. In all three models, we included petri dish identity as a random intercept effect because ants from the same dish are non-independent. Additionally, a random intercept effect was included for each ant as we observed the same individuals twice (before and after unpacking).

### Chemical bioassay

We determined whether ants detect infected pupae through potential changes in the pupae’s cuticular chemical profile. We established internal infections in pupae by exposing them to the pathogen and leaving them for 3 d in isolation. In pilot studies, approx. 50% of these pupae were then unpacked within 4 h of being introduced to ants. After 3 d, pupae were washed for 2.5 min in 300 μl of either pentane solvent to reduce the abundance of all CHCs present on the pupae (*n* = 28), or in autoclaved water as a handling control (*n* = 28). After washing, pupae were allowed to air dry on sterile filter paper. Additionally, non-washed pupae were used as a positive control (*n* = 30). Pupae were placed individually with a pair of ants in petri dishes and observed for unpacking for 4 h. We used GC–MS (see below for methodology) to confirm that washing was effective at removing cuticular compounds, by comparing the total amount of chemicals present on pupae washed in pentane to non- and water-washed pupae (*n* = 8 per treatment; Fig S4). The number of pupae unpacked between the different treatments was analysed using a logistic regression (Fig 2A). As several researchers helped to wash the pupae, we included a random intercept for each person to control for any potential handling effects. Additionally, the experiment was run in two blocks on separate days, so we included a random intercept for each block to generalise beyond any potential differences between runs. The total peak area from the GC–MS analysis was compared between treatments using a KW test with post hoc comparisons.

### Chemical analysis of pupal hydrocarbon patterns

To confirm that infected pupae had chemical profiles that are different from pathogen-exposed cocooned and control pupae, we exposed pupae to the pathogen or a sham control. Pupae were then isolated for 3 d to establish infections in the pathogen-exposed treatment (as above). Following isolation, pupae were individually placed with ants and observed for unpacking for 4 h. Unpacked pupae were immediately frozen at – 80 °C with the removed cocoons (*n* = 13) and we also froze cocooned pathogen-exposed pupa that had not yet been unpacked (*n* =10). Furthermore, we froze a pair of control pupae, of which one was cocooned (*n* = 12), whilst the other was first experimentally unpacked (to test if the cocoon affects cuticular compound extraction; *n* =12). Cuticular chemicals were extracted from individual pupae and their cocoons in glass vials (Supelco; 1.8 ml) containing 100 μl n-pentane solvent for 5 min under gentle agitation. The vials were then centrifuged at 3000 rpm for 1 min to spin down any fungal conidiospores that might be remaining, and 80 μl of the supernatant was transferred to fresh vials with 200 μl glass inserts and sealed with Teflon faced silicon septa (both Supelco). The pentane solvent contained four internal standards relevant for our range of hydrocarbons (C_27_ – C_37_); n-Tetracosane, n-Triacontane, n-Dotriacontane and n-Hexatriacontane (Sigma Aldrich) at 0.5 μg/ml concentration, all fully deuterated to enable spectral traceability and separation of internal standards from ant-derived substances. We ran extracts from the different groups in a randomised manner, intermingled with blank runs containing only pentane, and negative controls containing the pentane plus internal standards (to exclude contaminants emerging e.g. from column bleeding), on the day of extraction, using GC–MS (Agilent Technologies; GC7890 coupled to MS5975C).

A liner with one restriction ring filled with borosilicate wool (Joint Analytical Systems) was installed in the programmed temperature vaporisation (PTV) injection port of the GC, which was pre-cooled to -20 °C and set to solvent vent mode. 50 μl of the sample extractions were injected automatically into the PTV port at 40 μl/s using an autosampler (CTC Analytics, PAL COMBI-xt; Axel Semrau, CHRONOS 4.2 software) equipped with a 100 μl syringe (Hamilton). Immediately after injection, the PTV port was ramped to 300 °C at 450 °C/min, and the sample transferred to the column (DB-5ms; 30 m × 0.25 mm, 0.25 μm film thickness) at a flow of 1 ml/min. The oven temperature program was held at 35 °C for 4.5 min, then ramped to 325 °C at 20 °C/min, and held at this temperature for 11 min. Helium was used as the carrier gas at a constant flow rate of 3 ml/min. For all samples, the MS transfer line was set to 325 °C, and the MS operated in electron ionisation mode (70 eV; ion source 230 °C; quadrupole 150 °C, mass scan range 35-600 amu, with a detection threshold of 150). Data acquisition was carried out using MassHunter Workstation, Data Acquisition software B.07.01 (Agilent Technologies).

Analytes were detected by applying deconvolution algorithms to the total ion chromatograms of the samples (MassHunter Workstation, Qualitative Analysis B.07.00). Compound identification (Table S1) was performed via manual interpretation using retention indices and spectral information, and the comparison of mass spectra to the Wiley 9^th^ edition/NIST 11 combined mass spectral database (National Institute of Standards and Technologies). As the molecular ion was not detectable for all analytes based on electronic ionisation, we in addition performed chemical ionisation on pools of 20 pupae in 100 μl n-pentane solvent with 0.5 μg/ml internal standards. The higher extract concentration was needed to counteract the loss in ionisation efficiency in chemical ionisation mode. A specialised chemical ionisation source with methane as the reagent gas was used with the MS, while the chromatographic method was the same as in electronic ionisation mode. Use of external standards (C_7_-C_40_ saturated alkane mixture [Sigma Aldrich]) enabled traceability of all peaks, and thus comparison to runs of single pupae extracts made in electronic ionisation mode. Modified Kovats retention indices for the peaks in question were calculated based on those standards. To further aid identification, we separated the substances based on polarity using solid phase extraction fractionation. For this purpose, pools of 20 pupae were extracted in 500 μl n-pentane containing 0.2 μg/ml internal standard, and separated on unmodified silica cartridges (Chromabond ^®^ SiOH, 1ml, 100 mg) based on polarity. Prior to use, the cartridges were conditioned with 1 ml dichloromethane followed by 1 ml n-pentane. The entire extraction volume was loaded onto the silica and the eluent (fraction 1, highly apolar phase) collected. A wash with 1 ml pure n-pentane was added to fraction 1. Fraction 2 contained all substances washed off the silica with 1 ml 25 % dichloromethane in n-pentane, and finally a pure wash with 1 ml dichloromethane eluted all remaining substances (fraction 3). The polarity thus increased from fraction 1 through 3, but no polar substances were found. All fractions were dried under a gentle nitrogen stream and re-suspended in 70 μl n-pentane followed by vigorous vortexing for 45 s. GC–MS analysis of all fractions was performed in electronic ionisation mode under the same chromatographic conditions as before.

To quantify the relative abundances of all compounds found on each pupa, analyte-characteristic quantifier and qualifier ions were used to establish a method enabling automatized quantification of their integrated peak area relative to the peak area of the closest internal standard. For each analyte, the relative peak area was normalised, i.e. divided by the total sum of all relative peak areas of one pupa, to standardise all pupa samples. Only analytes, which normalised peak area contributed more than 0.05% of the total peak area, were included in the statistical analysis. We compared the chemical profiles of the pupae using a perMANOVA analysis of the Mahalanobis dissimilarities between pupae, with post hoc perMANOVA comparisons. Since there was no difference between cocooned and unpacked control pupae we combined them into a single control group for the final analysis (perMANOVA: *F* = 1.09, df = 23, *P* = 0.1). We also performed a discriminant analysis of principle components (Fig 2B) to characterise the differences between the pupal treatments [88,89]. To identify the compounds that differ between treatments, we performed a conditional random forest classification (*n* trees = 500, *n* variables per split = 4) [88,90,91]. Random forest identified 9 compounds that were important in classifying the treatment group, of which 8 were significant when analysed using separate KW tests (results for significant compounds in Table S2). We followed up the KW tests with individual post hoc comparisons for each significant compound (Fig 2C-F, post hoc comparisons in Table S2).

### Effect of destructive disinfection on pathogen replication

To test if destructive disinfection prevents *Metarhizium* from successfully replicating, we kept single pathogen-exposed pupae in petri dishes containing groups of 3 or 8 ants. This allowed us to assess how group size affects the likelihood of fungal inhibition. For the following 10 d, we observed the pupae for unpacking. When a pupa was unpacked, we left it with the ants for a further 1 or 5 d so that they could perform destructive disinfection. This allowed us to assess how the duration of destructive disinfection affects the likelihood of fungal inhibition. The destructively disinfected pupae were then removed and placed into petri dishes on damp filter paper at 23 °C (8 ants 1 d and 5 d, *n* = 22 pupae each; 3 ants 1 and 5 d, *n* = 18 pupae each). We did not surface sterilise the pupae as this might have interfered with the destructive disinfection the ants had performed. Removed pupae were observed daily for *Metarhizium* sporulation for 30 d. To determine how many pupae sporulate in the absence of destructive disinfection, we kept pathogen-exposed pupae without ants as a control and recorded the number that sporulated for 30 d (*n* = 25). We compared the number of pupae that sporulated after 1 d, 5 d and in the absence of ants using logistic regressions and Tukey post hoc comparisons, separately for the two ant group sizes (Fig 3A, Fig S6).

### *In vitro* investigation of destructive disinfection

We examined the individual effects of unpacking, biting and poison application on destructive disinfection by performing these behaviours *in vitro*. Pathogen-exposed pupae were initially kept with ants so that they could perform sanitary care. After 3 d, we removed the pupae and split them up into three groups: (i) pupae that we left cocooned, (ii) experimentally unpacked and (iii) experimentally unpacked and bitten. We simulated the damage the ants achieve through biting by damaging the pupal cuticle and removing their limbs with forceps. The pupae were then treated with either synthetic ant poison (60% formic acid and 2% acetic acid, in water; applied at a dose equivalent to what ants apply during destructive disinfection; Fig S8) or autoclaved distilled water as a control, using pressurised spray bottles (Lacor) to evenly coat the pupae in liquid. The pupae were allowed to air dry for 5 min before being rolled over and sprayed again and allowed to dry a further 5 min. All pupae were then placed into separate petri dishes and monitored daily for *Metarhizium* sporulation (cocooned + poison, *n* = 24; unpacked + poison + biting, *n* = 24; all other treatments, *n* = 25). The number of pupae sporulating was analysed using a logistic regression with Firth’s penalised likelihood, which offers a solution to the monotone likelihood caused by the complete absence of sporulation in one of the groups (R package ‘brglm’ [92]). Pupal manipulation (cocooned/unpacked only/unpacked and bitten), chemical treatment (water or poison) and their interaction were included as main effects (Fig 3B, Fig S7). We followed up this analysis with Tukey post hoc comparisons (Table S3).

### Disease transmission from infectious and destructively disinfected pupae

We tested the impact of destructive disinfection on disease transmission within groups of ants by keeping them with sporulating pupae or pupae that had been destructively disinfected. Infections were established in pupae (as above) and half were allowed to sporulate (*n* = 11), whilst the other half were experimentally destructively disinfected (as above; *n* = 11). Pupae were then kept individually with groups of 5 ants in mini-nests (cylindrical containers [Ø = 90 mm] with a second, smaller chamber covered in red foil [Ø = 33 mm]). Ant mortality was monitored daily for 30 d. Dead ants were removed, surface sterilised and observed for *Metarhizium* sporulation. The number of ants dying from *Metarhizium* infections in each treatment was compared using a logistic regression (Fig 4A). Mini-nest identity was included as a random intercept effect as ants from the same group are non-independent.

## Acknowledgments

We thank L. Lovicar for producing SEMs, B. Leyrer for help with chemical analysis, B. Milutinović and M. Bračić for assistance with the chemical bioassay and the *Social Immunity* group at IST Austria for ant collections and comments on earlier drafts of the manuscript. Finally, we are grateful to M. Sixt, D. Siekhaus and J. J. Boomsma for discussion of the project throughout.

## References

1. Schmid-Hempel P. Evolutionary parasitology: the integrated study of infections, immunology, ecology, and genetics. New York: Oxford University Press; 2011.

2. Nunn CL, Altizer S. Infectious diseases in primates: behavior, ecology and evolution. New York: Oxford University Press; 2006.

3. Schmid-Hempel P. Parasites and their social hosts. Trends Parasitol. 2017; doi:10.1016/j.pt.2017.01.003

4. Alexander RD. The evolution of social behavior. Annu Rev Ecol Syst. 1974; 5: 325–383. doi:10.1146/annurev.es.05.110174.001545

5. Hamilton WD. Kinship, recognition, disease, and intelligence: constraints of social evolution. In: Ito Y, Brown J, Kikkawa J, editors. Animal societies: theories and facts. Tokyo: Japan Sci. Soc. Press; 1987. pp. 81–102.

6. Ezenwa VO, Ghai RR, McKay AF, Williams AE. Group living and pathogen infection revisited. Curr Opin Behav Sci. 2016; 12: 66–72. doi:10.1016/j.cobeha.2016.09.006

7. Queller DC, Strassmann JE. Eusociality. Curr Biol. 2003; 13: R861–R863. doi:10.1016/j.cub.2003.10.043

8. Bourke AFG. Principles of social evolution. Oxford: Oxford University Press; 2011.

9. West SA, Fisher RM, Gardner A, Toby Kiers E. Major evolutionary transitions in individuality. Proc Natl Acad Sci. 2015; 112: 10112–10119. doi:10.1073/pnas.1421402112

10. Wheeler WM. The ant-colony as an organism. J Morphol. 1911; 22: 307–325. doi:10.1002/jmor.1050220206

11. Boomsma JJ, Gawne R. Superorganismality and caste differentiation as points of no return: how the major evolutionary transitions were lost in translation. Biol Rev. 2017;Forthcoming.

12. Cremer S, Armitage SAO, Schmid-Hempel P. Social immunity. Curr Biol. 2007; 17: 693–702. doi:10.1016/j.cub.2007.06.008

13. Shorter JR, Rueppell O. A review on self-destructive defense behaviors in social insects. Insectes Soc. 2011; 59: 1–10. doi:10.1007/s00040-011-0210-x

14. Stroeymeyt N, Casillas-Pérez B, Cremer S. Organisational immunity in social insects. Curr Opin Insect Sci. 2014; 5: 1–15. doi:10.1016/j.cois.2014.09.001

15. Wilson-Rich N, Spivak M, Fefferman NH, Starks PT. Genetic, individual, and group facilitation of disease resistance in insect societies. Annu Rev Entomol. 2009; 54: 405–23. doi:10.1146/annurev.ento.53.103106.093301

16. Meunier J. Social immunity and the evolution of group living in insects. Philos Trans R Soc B Biol Sci. 2015; 370: 20140102. doi:10.1098/rstb.2014.0102

17. Cremer S, Sixt M. Analogies in the evolution of individual and social immunity. Philos Trans R Soc London B Biol Sci. 2009; 364: 129–42. doi:10.1098/rstb.2008.0166

18. Tranter C, LeFevre L, Evison SEF, Hughes WOH. Threat detection: contextual recognition and response to parasites by ants. Behav Ecol. 2015; 26: 396–405. doi:10.1093/beheco/aru203

19. Rosengaus R, Traniello JFA. Disease susceptibility and the adaptive nature of colony demography in the dampwood termite *Zootermopsis angusticollis*. Behav Ecol Sociobiol. 2001; 50: 546–556. doi:10.1007/s002650100394

20. Reber A, Purcell J, Buechel SD, Buri P, Chapuisat M. The expression and impact of antifungal grooming in ants. J Evol Biol. 2011; 24: 954–64. doi:10.1111/j.1420-9101.2011.02230.x

21. Graystock P, Hughes WOH. Disease resistance in a weaver ant, *Polyrhachis dives*, and the role of antibiotic-producing glands. Behav Ecol Sociobiol. 2011; 65: 2319–2327. doi:10.1007/s00265-011-1242-y

22. Tragust S, Mitteregger B, Barone V, Konrad M, Ugelvig L V, Cremer S. Ants disinfect fungus-exposed brood by oral uptake and spread of their poison. Curr Biol. 2013; 23: 76–82. doi:10.1016/j.cub.2012.11.034

23. Hughes WOH, Eilenberg J, Boomsma JJ. Trade-offs in group living: transmission and disease resistance in leaf-cutting ants. Proc Biol Sci B Biol Sci. 2002; 269: 1811–1819. doi:10.1098/rspb.2002.2113

24. Loreto RG, Hughes DP. Disease in the society: infectious cadavers result in collapse of ant sub-colonies. PLoS One. 2016; 11: e0160820. doi:10.1371/journal.pone.0160820

25. Deacon JW. Fungal Biology. Malden: Blackwell Publishing; 2006.

26. Ugelvig L V, Kronauer DJC, Schrempf A, Heinze J, Cremer S. Rapid anti-pathogen response in ant societies relies on high genetic diversity. Proc R Soc B. 2010; 277: 2821–2828. doi:10.1098/rspb.2010.0644

27. Tragust S, Ugelvig L V, Chapuisat M, Heinze J, Cremer S. Pupal cocoons affect sanitary brood care and limit fungal infections in ant colonies. BMC Evol Biol. 2013; 13: 225. doi:10.1186/1471-2148-13-225

28. Okuno M, Tsuji K, Sato H, Fujisaki K. Plasticity of grooming behavior against entomopathogenic fungus *Metarhizium anisopliae* in the ant *Lasius japonicus*. J Ethol. 2011; 30: 23–27. doi:10.1007/s10164-011-0285-x

29. Leonhardt SD, Menzel F, Nehring V, Schmitt T. Ecology and evolution of communication in social insects. Cell. 2016; 164: 1277–1287. doi:10.1016/j.cell.2016.01.035

30. Theraulaz G, Bonabeau E, Deneubourg J. Response threshold reinforcements and division of labour in insect societies. Proc R Soc B Biol Sci. 1998; 265: 327–332. doi:10.1098/rspb.1998.0299

31. Beshers SN, Fewell JH. Models of division of labor in social insects. Annu Rev Entomol. 2001; 46: 413–440. doi:10.1146/annurev.ento.46.1.413

32. Sharma KR, Enzmann BL, Schmidt Y, Moore D, Jones GR, Parker J, et al. Cuticular hydrocarbon pheromones for social behavior and their coding in the ant antenna. Cell Rep. 2015; 1–11. doi:10.1016/j.celrep.2015.07.031

33. Hefetz A, Blum MS. Biosynthesis of formic acid by the poison glands of formicine ants. Biochim Biophys Acta. 1978; 543: 484–496. doi:10.1016/0304- 4165(78)90303-3

34. Walch M, Dotiwala F, Mulik S, Thiery J, Kirchhausen T, Clayberger C, et al. Cytotoxic cells kill intracellular bacteria through granulysin-mediated delivery of granzymes. Cell. 2014;157: 1309–1323. doi:10.1016/j.cell.2014.03.062

35. Kägi D, Ledermann B, Bürki K, Seiler P, Odermatt B, Olsen KJ, et al. Cytotoxicity mediated by T cells and natural killer cells is greatly impaired in perforin-deficient mice. Nature. 1994; 369: 31–37. doi:10.1038/369031a0

36. Chowdhury D, Lieberman J. Death by a thousand cuts: granzyme pathways of programmed cell death. Annu Rev Immunol. 2008; 26: 389–420. doi:10.1146/annurev.immunol.26.021607.090404

37. Shore SL, Cromeans TL, Romano TJ. Immune destruction of virus-infected cells early in the infectious cycle. Nature. 1976; 262: 695–696. doi:10.1038/262695a0

38. Leclerc J-B, Detrain C. Ants detect but do not discriminate diseased workers within their nest. Sci Nat. 2016; 103: 70. doi:10.1007/s00114-016-1394-8

39. Bos N, Lefèvre T, Jensen AB, D’Ettorre P. Sick ants become unsociable. J Evol Biol. 2012; 25: 342–51. doi:10.1111/j.1420-9101.2011.02425.x

40. Diez L, Moquet L, Detrain C. Post-mortem changes in chemical profile and their influence on corpse removal in ants. J Chem Ecol. 2013; 39: 1424–1432. doi:10.1007/s10886-013-0365-1

41. Qiu H-L, Lu L-H, Shi Q-X, Tu C-C, Lin T, He Y-R. Differential necrophoric behaviour of the ant *Solenopsis invicta* towards fungal-infected corpses of workers and pupae. Bull Entomol Res. 2015; 105: 607–614. doi:10.1017/S0007485315000528

42. Wilson EO, Durlach NI, Roth LM. Chemical releaser of necrophoric behavior in ants. Psyche. 1958; 65: 108–114. doi:10.1155/1958/69391

43. Trabalon M, Plateaux L, Péru L, Bagnères A-G, Hartmann N. Modification of morphological characters and cuticular compounds in worker ants *Leptothorax nylanderi* induced by endoparasites *Anomotaenia brevis*. J Insect Physiol. 2000; 46: 169–178. doi:10.1016/S0022-1910(99)00113-4

44. Lin L, Fang W, Liao X, Wang F, Wei D, St. Leger RJ. The MrCYP52 cytochrome P450 monoxygenase gene of *Metarhizium robertsii* is important for utilizing insect epicuticular hydrocarbons. PLoS One. 2011; 6: e28984. doi:10.1371/journal.pone.0028984

45. Richard F-J, Aubert A, Grozinger C. Modulation of social interactions by immune stimulation in honey bee, *Apis mellifera*, workers. BMC Biol. 2008; 6: 50. doi:10.1186/1741-7007-6-50

46. Richard F-J, Holt HL, Grozinger CM. Effects of immunostimulation on social behavior, chemical communication and genome-wide gene expression in honey bee workers (*Apis mellifera*). BMC Genomics. 2012; 13: 558. doi:10.1186/1471-2164-13-558

47. Baracchi D, Fadda A, Turillazzi S. Evidence for antiseptic behaviour towards sick adult bees in honey bee colonies. J Insect Physiol. 2012; 58: 1589–1596. doi:10.1016/j.jinsphys.2012.09.014

48. Shakhar K, Shakhar G. Why do we feel sick when infected—can altruism play a role? PLoS Biol. 2015; 13: e1002276. doi:10.1371/journal.pbio.1002276

49. Ravichandran KS. Find-me and eat-me signals in apoptotic cell clearance: progress and conundrums. J Exp Med. 2010; 207: 1807–1817. doi:10.1084/jem.20101157

50. Grimsley C, Ravichandran KS. Cues for apoptotic cell engulfment: eat-me, don’t eat-me and come-get-me signals. Trends Cell Biol. 2003; 13: 648–656. doi:10.1016/j.tcb.2003.10.004

51. Tragust S. External immune defence in ant societies (Hymenoptera: Formicidae): the role of antimicrobial venom and metapleural gland secretion. Myrmecol news. 2016;23: 119–128.

52. Knaden M, Wehner R. Nest defense and conspecific enemy recognition in the desert ant *Cataglyphis fortis*. J Insect Behav. 2003; 16: 717–730. doi:10.1023/B:JOIR.0000007706.38674.73

53. Reber A, Chapuisat M. Diversity, prevalence and virulence of fungal entomopathogens in colonies of the ant *Formica selysi*. Insectes Soc. 2012; 59: 231–239. doi:10.1007/s00040-011-0209-3

54. Hughes WOH, Thomsen L, Eilenberg J, Boomsma JJ. Diversity of entomopathogenic fungi near leaf-cutting ant nests in a neotropical forest, with particular reference to *Metarhizium anisopliae* var. *anisopliae*. J Invertebr Pathol. 2004; 85: 46–53. doi:10.1016/j.jip.2003.12.005

55. Cremer S, Ugelvig L V, Drijfhout FP, Schlick-Steiner BC, Steiner FM, Seifert B, et al. The evolution of invasiveness in garden ants. PLoS One. 2008; 3: e3838. doi:10.1371/journal.pone.0003838

56. Keller S, Kessler P, Schweizer C. Distribution of insect pathogenic soil fungi in Switzerland with special reference to *Beauveria brongniartii* and *Metharhizium anisopliae*. BioControl. 2003; 48: 307–319. doi:10.1023/A:1023646207455

57. Roberts DW, St. Leger RJ. *Metarhizium* spp., cosmopolitan insect-pathogenic fungi: mycological aspects. Adv Appl Microbiol. 2004. pp. 1–70. doi:10.1016/S0065-2164(04)54001-7

58. Shimazu M. *Metarhizium cylindrosporae* Chen et Guo (Deuteromycotina: Hyphomycetes), the causative agent of an epizootic on *Graptopsaltria nigrofuscata* Motchulski (Homoptera: Cicadiae). Appl Entomol Zool. 1989; 24: 430–434. doi:10.1303/aez.24.430

59. Lomer CJ, Prior C, Kooyman C. Development of *Metarhizium* spp. for the control of grasshoppers and locusts. Mem Entomol Soc Canada. 1997; 129: 265–286. doi:10.4039/entm129171265-1

60. Hughes DP, Araújo J, Loreto R, Quevillon L, de Bekker C, Evans HC. From so simple a beginning: the evolution of behavioral manipulation by fungi. Adv Genet. 2016; 94: 1–33. doi:10.1016/bs.adgen.2016.01.004

61. Loreto RG, Elliot SL, Freitas MLR, Pereira TM, Hughes DP. Long-term disease dynamics for a specialized parasite of ant societies: a field study. PLoS One. 2014; 9: e103516. doi:10.1371/journal.pone.0103516

62. Malagocka J, Jensen AB, Eilenberg J. *Pandora formicae*, a specialist ant pathogenic fungus: new insights into biology and taxonomy. J Invertebr Pathol. 2017; 143: 108–114. doi:10.1016/j.jip.2016.12.007

63. Agosta SJ, Janz N, Brooks DR. How specialists can be generalists: resolving the “parasite paradox” and implications for emerging infectious disease. Zoologia. 2010; 27: 151–162. doi:10.1590/S1984-46702010000200001

64. Verza SS, Diniz EA, Chiarelli MF, Mussury RM, Bueno OC. Waste of leaf-cutting ants: disposal, nest structure, and abiotic soil factors around internal waste chambers. Acta Ethol.; 2017; doi:10.1007/s10211-017-0255-6

65. Hart AG, Ratnieks FLW. Waste management in the leaf-cutting ant *Atta colombica*. Behav Ecol. 2002; 13: 224–231. doi:10.1093/beheco/13.2.224

66. Farji-Brener AG, Elizalde L, Fernández-Marín H, Amador-Vargas S. Social life and sanitary risks: evolutionary and current ecological conditions determine waste management in leaf-cutting ants. Proc R Soc B Biol Sci. 2016; 283: 20160625. doi:10.1098/rspb.2016.0625

67. Spivak M, REuter GS. Resistance to American foulbrood disease by honey bee colonies *Apis mellifera* bred for hygienic behavior. Apidologie. 2001; 32: 555–565. doi:10.1051/apido:2001103

68. Chouvenc T, Su N-Y. When subterranean termites challenge the rules of fungal epizootics. PLoS One. 2012; 7: e34484. doi:10.1371/journal.pone.0034484

69. Chouvenc T, Su N-Y, Robert A. Inhibition of *Metarhizium anisopliae* in the alimentary tract of the eastern subterranean termite *Reticulitermes flavipes*. J Invertebr Pathol. 2009; 101: 130–6. doi:10.1016/j.jip.2009.04.005

70. Rosengaus RB, Schultheis KF, Yalonetskaya A, Bulmer MS, DuComb WS, Benson RW, et al. Symbiont-derived β-1,3-glucanases in a social insect: mutualism beyond nutrition. Front Microbiol. 2014; 5: 1–11. doi:10.3389/fmicb.2014.00607

71. Rosengaus R, Guldin M, Traniello JFA. Inhibitory effect of termite fecal pellets on fungal spore germination. J Chem Ecol. 1998; 24: 1697–1706. doi:10.1023/A:1020872729671

72. Rosengaus RB, Traniello JFA, Bulmer MS. Ecology, behavior and evolution of disease resistance in termites. In: Bignell DE, Roisin Y, Lo N, editors. Biology of termites: a modern synthesis. New York: Springer; 2011. pp. 165–191.

73. R Core Team. R: A language and environment for statistical computing. Vienna, Austria: R Foundation for Statistical Computing; 2012.

74. Bolker BM, Brooks ME, Clark CJ, Geange SW, Poulsen JR, Stevens MHH, et al. Generalized linear mixed models: a practical guide for ecology and evolution. Trends Ecol Evol. 2009; 24: 127–35. doi:10.1016/j.tree.2008.10.008

75. Benjamini Y, Hochberg Y. Controlling the false discovery rate: a practical and powerful approach to multiple testing. J R Stat Soc. 1995; 57: 289–300. doi:10.2307/2346101

76. García L V. Escaping the Bonferroni iron claw in ecological studies. Oikos. 2004; 105: 657–663. doi:10.1111/j.0030-1299.2004.13046.x

77. Bates D, Maechler M, Bolker B, Walker S. lme4: linear mixed-effects models using Eigen and S4. R Foundation for Statistical Computing; 2014.

78. Nieuwenhuis R, Pelzer B, te Grotenhuis M. influence.ME: tools for detecting influential data in mixed effects models. R J. 2012;4: 38–47.

79. Kuznetsova, Alexandra, Brockhoff PB, Christensen RHB. lmerTest: tests in linear mixed effects models. R Foundation for Statistical Computing; 2015.

80. Ploner M, Heinze G. coxphf: Cox regression with Firth’s penalized likelihood. R Foundation for Statistical Computing; 2015.

81. de Mendiburu F. agricolae: statistical procedures for agricultural research. R Foundation for Statistical Computing; 2016.

82. Oksanen J, Blanchet FG, Kindt R, Legendre P, Minchin PR, O’Hara B, et al. vegan: community ecology package. R Foundation for Statistical Computing; 2016.

83. Bretz F, Hothorn T, Westfall P. Multiple comparisons using R. Boca Ration: CRC Press; 2011.

84. Wickham H. ggplot2: elegant graphics for data analysis. New York: Springer; 2009.

85. Lacey LA, Brooks WM. Initial handling and diagnosis of diseased invertebrates. In: Lacey LA, editor. Manual of techniques in invertebrate pathology. San Diego: Academic Press; 1997. p. 5.

86. Blumstein DT, Daniel JC. Quantifying behavior the JWatcher way. Sunderland: Sinauer Associates; 2007.

87. Harrison XA. Using observation-level random effects to model overdispersion in count data in ecology and evolution. PeerJ. 2014; 2: e616. doi:10.7717/peerj.616

88. De Moraes CM, Stanczyk NM, Betz HS, Pulido H, Sim DG, Read AF, et al. Malaria-induced changes in host odors enhance mosquito attraction. Proc Natl Acad Sci. 2014; 111: 11079–11084. doi:10.1073/pnas.1405617111

89. Jombart T, Devillard S, Balloux F. Discriminant analysis of principal components: a new method for the analysis of genetically structured populations. BMC Genet. 2010; 11: 94. doi:10.1186/1471-2156-11-94

90. Strobl C, Hothorn T, Zeileis A. Party on! R J. 2009;1: 14–17.

91. Strobl C, Malley J, Tutz G. An introduction to recursive partitioning: rationale, application, and characteristics of classification and regression trees, bagging, and random forests. Psychol Methods. 2009; 14: 323–348. doi:10.1037/a0016973

92. Kosmidis I. brglm: Bias reduction in binomial-response generalized linear models. R Foundation for Statistical Computing; 2013.

